# Synaptic anchoring of the endoplasmic reticulum depends on myosin V and caldendrin activity

**DOI:** 10.1101/2020.08.14.250746

**Authors:** Anja Konietzny, Jasper Grendel, Nathalie Hertrich, Dick H. W. Dekkers, Jeroen A. A. Demmers, Marina Mikhaylova

**Affiliations:** AG Optobiology, Humboldt Universität zu Berlin, Institute of Biology, Invalidenstraße 42, D-10115 Berlin, Germany; Guest Group “Neuronal Protein Transport”, Center for Molecular Neurobiology, ZMNH, University Medical Center Hamburg-Eppendorf, 20251 Hamburg, Germany; Center for Proteomics, Erasmus MC, 3000 CA Rotterdam, The Netherlands

**Keywords:** caldendrin, endoplasmic reticulum, myosin, spine apparatus, synapse

## Abstract

Excitatory synapses of principal hippocampal neurons are frequently located on dendritic spines. The dynamic strengthening or weakening of individual inputs results in a great structural and molecular diversity of dendritic spines. Active spines with large Ca^2+^ transients are frequently invaded by a single protrusion from the endoplasmic reticulum (ER), which is dynamically transported into and out of spines by the actin-based motor myosin V. An increase in synaptic strength often correlates with stable anchoring of the ER, followed by the formation of the spine apparatus organelle. Here we show that synaptic ER stabilization depends on the interplay of two Ca^2+^-binding proteins: calmodulin serves as a light chain of myosin V and activates the motor function, whereas caldendrin acts as an inhibitor which transforms myosin into a stationary F-actin tether. Together, they provide a Ca^2+^-sensing module for fine-tuning myosin V activity and thereby regulate the formation of the spine apparatus in a subset of active dendritic spines.

## Introduction

The dynamic plasticity of neuronal synapses is believed to be the basis of adaptive neuronal circuit formation, which underlies complex brain functions such as learning and memory. The majority of excitatory synapses on neuronal dendrites are located on small membranous protrusions called dendritic spines. Spines are highly dynamic structures that can undergo extensive morphological changes, appear, disappear or become stabilized over long periods of time (weeks and months), depending on the type of synaptic input they receive (Wiegert *et al*, 2018). The structural plasticity of dendritic spines depends on the tight regulation of spinous Ca^2+^-signaling, and on local, activity-dependent protein and membrane turnover. Dendritic secretory trafficking organelles play an important role in this regard (Mikhaylova *et al*, 2016; Bowen *et al*, 2017; Goo *et al*, 2017; Padamsey *et al*, 2017; Yu *et al*, 2006; Kennedy & Hanus, 2019; Hanus *et al*, 2014; Holbro *et al*, 2009; Bommel *et al*, 2019).

How does neuronal activity regulate input-specific synaptic transport and organelle localization? Fast, long-distance cargo trafficking inside the dendrite is carried out by microtubule (MT)-based kinesin and dynein motor proteins, whereas locally confined transport and anchoring relies on F-actin-based myosin motors (Konietzny *et al*, 2017; Bommel *et al*, 2019). Unconventional myosins, and especially the myosin V (myoV) motor family can take over MT-based transport and redirect cargo into actin-rich areas such as dendritic spines (Kapitein *et al*, 2013; Bommel *et al*, 2019; Wang *et al*, 2008; Esteves da Silva *et al*, 2015). Two paralogs, myoVa and Vb, are highly expressed in the brain and are very similar in terms of their regulation and function (Wang *et al*, 2008; Hammer & Wagner, 2013). They were shown to mediate the synaptic transport of various cargoes, including recycling endosomes, endo-lysosomes, smooth endoplasmic reticulum (SER) and mRNA (Esteves da Silva *et al*, 2015; Kneussel & Wagner, 2013; van Bommel et al., 2019; Perez-Alvarez *et al*, 2020). Of note, in addition to its role as a processive motor, myoV can also serve as an organelle tether (Wagner *et al*, 2011; Maschi *et al*, 2018, Bommel *et al*, 2019). This could be especially important for tagging and diversification of dendritic spines.

In a very recent study on hippocampal neurons, myoV was shown to be responsible for the insertion of SER into highly active dendritic spines, which changed their plastic properties compared to SER-negative spines (Perez-Alvarez *et al*, 2020). Similarly, in an earlier study we described the involvement of myoVa in the spine localization of a more elaborate ER-derived organelle called the spine apparatus, which is a hallmark of large, stable spines (Konietzny *et al*, 2019). Importantly, while the motor function of myoV can be activated by cargo binding, it also depends on intracellular Ca^2+^ levels. Thus, regulated switching between processive cargo transport and stationary cargo anchoring on actin filaments could fine-tune organelle localization at dendritic spines. Together, this makes myoV an interesting candidate to mediate non-random, activity-dependent targeting of dendritic spines (Trybus *et al*, 2007).

What are the mechanisms that transduce changes in local Ca^2+^ concentrations to myoV activity? The mammalian myoV motor protein consists of two heavy chains, each of which contains six consensus binding sites (IQ motifs) for the ubiquitously expressed Ca^2+^-binding protein calmodulin (CaM). CaM binding necessary for stabilizing the lever arm, which translates the force generated by ATP hydrolysis in the motor domain into a forward movement. Especially CaM bound to the first IQ motif (IQ1-CaM) has been implied in the Ca^2+^-dependent activation of myoV activity *in vitro* (Krementsov *et al*, 2004; Lu *et al*, 2012; Shen *et al*, 2016; Trybus *et al*, 2007). However, is it still an open question how myoV-propelled cargo in dendritic spines can undergo a switch from active transport to local anchoring for longer time periods to ensure that SER or other membrane organelles can perform their synapse-related functions.

Here, we explored the mechanism of Ca^2+^-dependent regulation of myoV and its implications for myoV-dependent organelle transport in dendritic spines. We discovered that another brain-specific calmodulin-related Ca^2+^-binding protein, caldendrin, is a new Ca^2+^-dependent interactor of myoV. Caldendrin adopts a folded conformation in which the carboxy-terminal domain masks an interface in the amino-terminus for protein-protein interaction, which is rapidly released by Ca^2+^ binding (Mikhaylova et al., 2019). It exhibits similar Ca^2+^-binding kinetics to CaM and requires low micromolar Ca^2+^ concentrations to become activated (Reddy *et al*, 2014). In contrast to the diffusely distributed CaM, caldendrin is specifically enriched in a subset of large dendritic spines, which already pre-allocates its function to a selected spine population (Laube *et al*, 2002; Dieterich *et al*, 2008). Accordingly, caldendrin is implicated in various activity-dependent pathways, including the stabilization of F-actin in dendritic spines and inositol-3-phosphate receptor (IP_3_R) signalling which are important for synaptic plasticity (Mikhaylova *et al*, 2018; Li *et al*, 2013).

In the present study, mass spectrometry screening combined with protein-protein binding assays revealed a direct interaction of caldendrin with a confined region of myoV containing the IQ1 motif. Using *in vitro* reconstitution and inducible motor-cargo assays in live cells, we demonstrate that caldendrin acts as an inhibitor of myosin processivity without interfering with its F-actin association. These findings were corroborated by an analysis of SER-spine-entry-dynamics in hippocampal neurons in the presence or absence of caldendrin. Our findings suggest that caldendrin acts as a “lock” that transitions myoV function from processive movement to stationary anchoring. In this model, the Ca^2+^-dependent interplay between CaM and caldendrin is a way to fine-tune myoV-dependent SER positioning, which is important for the formation of the spine apparatus in dendritic spines.

## Results

### Caldendrin specifically interacts with a confined region of myoV containing the IQ1 motif

In our previous work on the caldendrin protein, we described its ability to undergo conformational change upon Ca^2+^ binding, its selective enrichment in a subset of large dendritic spines and its role in stabilizing actin filaments through interaction with cortactin (Reddy *et al*, 2014; Mikhaylova *et al*, 2018; Dieterich *et al*, 2008). These characteristics place caldendrin at an ideal position to mediate the consolidation of synaptic growth after a plasticity event.

To gain further insight into the Ca^2+^-dependent interactome of caldendrin, we performed a biotin-streptavidin pull-down of eukaryotically produced biotinylated GFP-caldendrin (bio-GFP-cald) or bio-GFP as a control from mouse brain lysate in the presence or absence of Ca^2+^ (Fig EV1A). Mass spectrometry (MS) analysis identified a broad network of putative brain-specific caldendrin binding partners (Table EV1). The obtained list of candidates was curated with a focus on the Ca^2+^-dependency of the interaction (Fig EV1B). We found that one class of proteins, myosins, were especially enriched in the caldendrin pull down in the presence of Ca^2+^. We hypothesized that, since the C-terminal domain of caldendrin is structurally very similar to calmodulin (CaM), and myosins contain known CaM binding sites (IQ motifs), those might mediate the caldendrin-myosin interaction (Fig EV1B,C). Of note, several other putative interactors identified in the MS also contain IQ motifs according to their UniProt annotation (Fig EV1B).

We decided to focus on one specific myosin, myoV, as it is was shown to be a driver of cargo-transport inside dendritic spines and as such important for spine plasticity and stability (Esteves da Silva *et al*, 2015; Yoshii *et al*, 2013; Perez-Alvarez *et al*, 2020).

To verify the MS results and to further characterize the caldendrin-myoV interaction, we employed a co-immunoprecipitation assay. Full-length, GFP-tagged myosin Va (GFP-myoVa fl) was co-expressed in HEK293T cells with full-length, tagRFP-coupled caldendrin (fl cald-tagRFP). GFP-myoVa was immunoprecipitated on GFP-trap beads from HEK293T cell lysate, which had been either supplemented with CaCl_2_ or with the Ca^2+^ chelator EGTA, and the presence of caldendrin was analyzed using western blot. To elucidate whether the interaction with myoVa was mediated via the caldendrin N-(aa1-136) or C-terminus (aa137-298), two truncation constructs were tested. We found that both fl caldendrin and the caldendrin N-terminus, but not the C-terminus, interacted with myoVa specifically in the presence of Ca^2+^ (Fig 1A). These results were surprising, as we would have expected the EF-hand containing C-terminal domain of caldendrin to mediate the interaction, due to its similarity to CaM and its ability to sense Ca^2+^. This suggests that the Ca^2+^-sensitivity of this interaction has to rely on the presence of another Ca^2+^ sensor.

**Figure 1:**
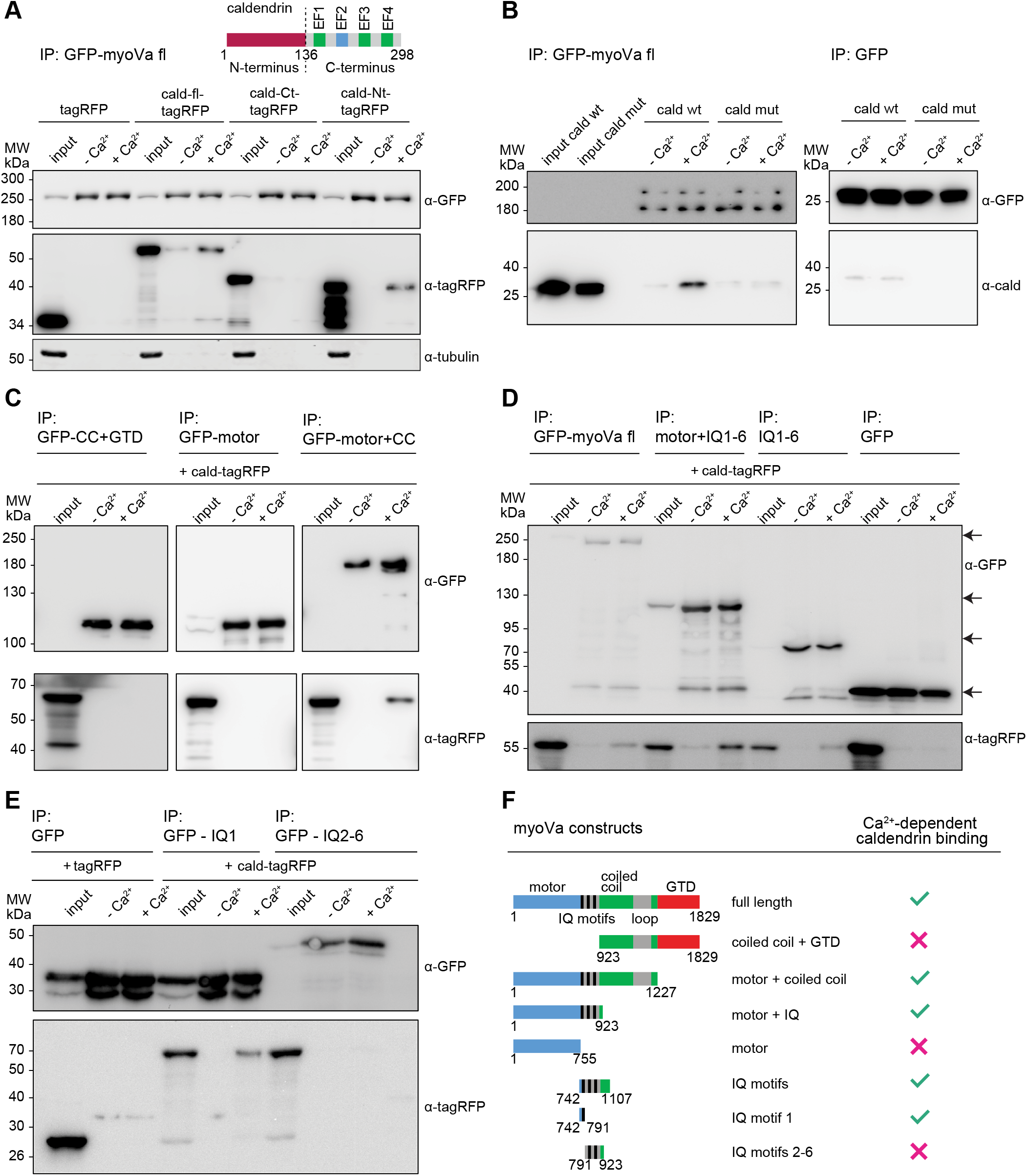
The calcium-binding protein caldendrin interacts with the first IQ motif of myosin Va in a Ca^2+^-dependent manner.

**A** Schematic showing overall domain organization of the caldendrin protein, including an unstructured N-terminal part, and four EF-hand binding motifs in the C-terminal part. Green boxes indicate functional EF-hands, blue indicates cryptic EF-hand. The dashed line indicates where the Nt and Ct constructs were separated. Co-immunoprecipitation experiments from extracts of HEK293T cells, co-expressing full-length GFP-myosin Va (myoVa fl) and indicated caldendrin-tagRFP (cald) constructs, show that caldendrin interacts with myoVa in a Ca^2+^-dependent manner via its N-terminus (Nt), and not via its C-terminus (Ct). Tubulin was used as a loading control for the input. MW = molecular weight marker.

**B** Pull down assay indicates that recombinant purified GFP-myoVa and untagged caldendrin (cald wt) directly bind to each other in a Ca^2+^-dependent manner. Note that there is no binding observed for a Ca^2+^-binding mutant of caldendrin (cald mut / Left panel). Right panel: neither wt nor mut caldendrin bind to GFP-coupled beads.

**C-E** Mapping of the binding region between myoVa and caldendrin using co-immunoprecipitation of cald-tagRFP and GFP-myoVa fragments expressed in HEK293 cells. Arrows in **D** indicate the size of the respective myosin fragment.

**F** Summary of the results from **C**– **E**. Caldendrin binds specifically to a fragment containing IQ-motif 1 (amino acids 742-791), but not to any other region of myoVa, in a Ca^2+^-dependent manner.

Further, to show that the myoVa-caldendrin interaction was indeed direct, and not mediated by an unknown adapter protein, we performed a pull-down assay using separately purified components. Here we used untagged caldendrin which was expressed and purified from *E. coli*. As an additional control we included a Ca^2+^-binding mutant (mut) of caldendrin that is locked in the closed and inactive conformation (Mikhaylova *et al*, 2018). Again, caldendrin showed a clear binding preference in the Ca^2+^ condition, whereas mut caldendrin showed little to no interaction in either condition (Fig 1B).

Next, we decided to map the binding region of caldendrin on the myoVa protein, by performing a co-immunoprecipitation assay using fl cald-tagRFP and various truncation constructs of GFP-myoVa (Fig 1C-E), summarized in Fig 1F. We found that caldendrin specifically interacts with a 49 amino-acid fragment (aa742-791, henceforth referred to as IQ1) containing part of the lever arm and the first IQ motif of myoVa in a Ca^2+^-dependent manner (Fig 1E,F).

### Caldendrin inhibits myosin V processivity *in vitro* possibly through competition with CaM

The results of binding interface mapping suggest that IQ1-CaM and caldendrin occupy very closely associated, or possibly overlapping binding sites on myoVa. Therefore, we next asked whether binding was competitive, or whether they could bind simultaneously. We expressed GFP-myoVa-IQ1 in HEK293T cells, coupled it to GFP-trap beads and added an excess of exogenous, recombinant CaM to make sure that all myoVa-IQ1 were fully occupied with CaM (Fig 2A,B, Fig EV2A). Additionally, we expressed tagRFP-cald in a separate batch of HEK293T cells and added the lysate to GFP-myoVa-IQ1-CaM-coupled beads (Fig 2A,B). Western blot analysis showed strong binding of caldendrin in the presence of Ca^2+^. It also revealed that the amount of CaM still bound to the beads after this incubation step was much lower in the presence of caldendrin and Ca^2+^, compared to the Ca^2+^-free condition (Fig 2B,C). Of note, we observed that in EGTA-supplemented cell lysate, CaM seemed to fully dissociate from myoVa-IQ1 (Fig 2B). This was unexpected, since according to literature, binding should be more stable in the absence of Ca^2+^ (Krementsov *et al*, 2004; Trybus *et al*, 2007). However, the fact that the amount of bound CaM seemed to be reduced in the Ca^2+^/caldendrin condition suggests that caldendrin, once it is bound to myoVa-IQ1, prevents CaM from re-binding, or that it might even displace CaM from its binding site.

**Figure 2:**
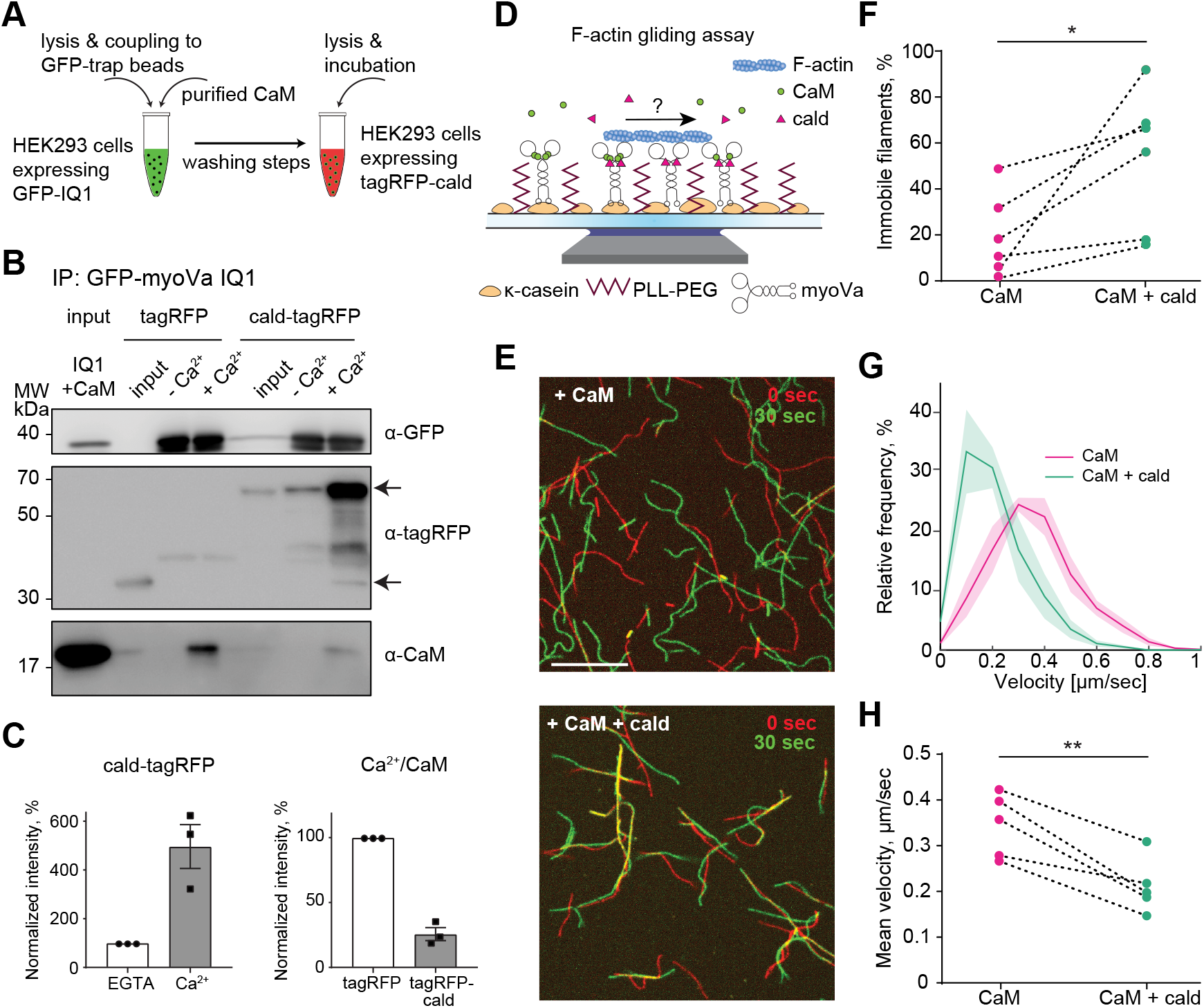
Caldendrin inhibits myosin Va processivity in in vitro F-actin gliding assay, possibly via replacement of calmodulin light chains at the IQ1 motif.

**A** Schematic of caldendrin and calmodulin competition assay. A construct containing the GFP-tagged first IQ-motif of myoVa (GFP-myoVa IQ1) was purified via GFP-trap beads from HEK293 cell lysate enriched with bacterially produced calmodulin (CaM). The CaM-associated construct (Input) was then incubated with lysate from HEK cells expressing either cald-tagRFP or tagRFP only.

**B** Western blot analysis of competition co-immunoprecipitation experiment in HEK293 cells. In the presence of Ca^2+^, caldendrin binds strongly to the IQ1 fragment (upper arrow), while tagRFP by itself does not bind (lower arrow). Detection using a calmodulin antibody indicated that Ca^2+^-dependent binding of caldendrin reduced the amount of CaM bound to the IQ1 motif relative to the control (tagRFP only). MW = molecular weight marker.

**C** Quantification of the normalized relative intensity of cald-tagRFP (left) and CaM bands (right) in western blot analysis of three independent experiments as shown in **B**. Shown is mean ± SEM. Left: Ca^2+^ greatly increases cald-tagRFP binding to GFP-myoVa-IQ1. Right: When Ca^2+^ is available, the simultaneous presence of cald-tagRFP leads to a reduced binding of CaM compared to the control (tagRFP).

**D** Schematic representation of *in vitro* F-actin gliding assay. CaM = Calmodulin, cald = caldendrin, PLL-PEG = Poly-L-lysine-polyethylene-glycol.

**E** Visualization of F-actin gliding caused by myoVa activity. The first (red) and last (green) frame of a 30 second time-lapse image stack were color-coded and overlaid to visualize movement. Yellow = actin filament did not move during 30 seconds of imaging. In the presence of only CaM (top), the majority of actin filaments are gliding. In the presence of both CaM and cald (bottom), a large number of filaments are immobile. Scale bar is 10 μm. Also see Movie EV2.

**F** Quantification of immobile filaments as shown in **E**. In the presence of caldendrin, the fraction of immobile actin filaments is significantly increased compared to CaM only (* p = 0.031, n=6 experiments, Wilcoxon matched-pairs signed rank test).

**G** Relative frequency distribution of recorded gliding velocities. In the presence of cald, the velocity distribution is shifted towards slower velocities compared to CaM only (mean ± SEM, n=5 experiments, 8 filaments for each experiment and condition).

**H** Quantification of F-actin gliding velocity. In the presence of cald, the velocity of F-actin gliding is significantly reduced compared to CaM only (mean ± SEM, ** p = 0.0052, paired t-test, n=5 experiments).

As Ca^2+^/CaM bound to IQ1 is known to play a role in myoVa activation (Trybus *et al*, 2007; Lu *et al*, 2012; Shen *et al*, 2016), we asked how caldendrin binding to myoVa might affect motor activity. To investigate this, we performed a reconstituted *in vitro* gliding assay that was designed as shown in Fig 2D. Full-length GFP-tagged myoVa was purified from HEK293 cells via a Twin-StrepTag (Fig EV2A,B) and attached to a glass surface in a flow chamber Alexa-568 labelled actin filaments were polymerized *in vitro* and introduced into the flow channel, where they could be observed to be “gliding” over the surface of myoVa motors. Untagged CaM and caldendrin were both purified from *E. coli* (Fig EV2C,D). To analyze the effect of caldendrin on the ability of myoVa to move F-actin, we took time-lapse image stacks of the gliding filaments and compared myoVa activity at 100 μM Ca^2+^ in the presence of CaM (30 μM) before and after supplementation with caldendrin (50 μM). Analyzed parameters were the percentage of “mobile” vs. “immobile” filaments in a given field of view (Fig 2E,F), and the velocity at which the myoVa motors were propelling the mobile actin filaments (Fig 2G,H). We found that the presence of caldendrin significantly increased the fraction of immobile filaments (from an average of 19.6 % to an average of 52.8 %; Fig 2F). At the same time, caldendrin strongly decreased the mobile filament velocity (from 344 nm/sec to 211 nm/sec; Fig 2G,H; Movie EV2). From this we concluded that caldendrin generally inhibits myoVa activity *in vitro*.

### Caldendrin does not affect the ability of myosin V to bind actin filaments

There are several ways in which caldendrin could inhibit myosin processivity. A simple explanation could be a conformational change which weakens the affinity of myoV for F-actin. However, this question could not be unambiguously answered by the gliding assay, as actin filaments can passively settle on the surface of the flow chamber without binding to myoV. Therefore, we investigated this possibility by first performing an F-actin pelleting assay using Ca^2+^-supplemented lysate from HEK293 cells that had been co-expressing either GFP-myoVa and cald-tagRFP, or GFP-myoVa and tagRFP only. In this assay, F-actin was stabilized during cell lysis by jasplakinolide, and subsequently pelleted via ultracentrifugation. F-actin-binding proteins co-sediment with F-actin, while soluble proteins (including monomeric G-actin) stay in the supernatant (Fig EV 3A). We found that tagRFP remained completely in the supernatant, whereas a fraction of cald-tagRFP had sedimented, presumably together with F-actin-bound myoVa or endogenous cortactin (Mikhaylova *et al*, 2018). In both conditions, myoVa was completely pelleted and did not seem to be affected by the presence of caldendrin. However, we could not exclude the possibility that GFP-myoVa was large enough to be “trapped” by the F-actin filaments and passively pulled into the pellet (Fig EV 3A).

Next, we reasoned that if caldendrin affects myoV affinity for F-actin, we should be able to observe a caldendrin-dependent change in myoV-dependent cargo trafficking. To investigate this we chose to use COS7 cells as a model system. First, since caldendrin binding to myoV requires the presence of Ca^2+^, we looked into COS7 endogenous Ca^2+^ dynamics by expressing the fluorescent Ca^2+^ sensor GCaMP7s (Dana *et al*, 2019). During 5 min live imaging periods, we observed that COS7 cells spontaneously produce large intracellular Ca^2+^ transients, making it plausible that in most cells that we looked at, caldendrin would have had a chance to bind to Ca^2+^, and thereby interact with myoV (Fig EV3B, Movie EV3.1). Next, we decided to employ an inducible cargo-motor coupling assay using peroxisomes, which are small and mostly immobile organelles. Peroxisomes can be coupled to different types of motor proteins via peroxisome-targeting protein sequences. The observed peroxisome motility then provides a readout of motor activity (Kapitein *et al*, 2013). Previous work has shown that, while kinesin-driven peroxisomes are being actively transported along microtubules in the cell periphery, additional recruitment of myoV is sufficient to stop or greatly reduce their motility due to myosin anchoring the peroxisome to F-actin (Kapitein *et al*, 2013). We hypothesized that, if caldendrin binding to myoV weakens its affinity for F-actin, myoV should be no longer able to stall kinesin-driven peroxisomes as effectively.

For this assay, we co-expressed three separate constructs: 1) a constitutively active, peroxisome-targeted (PEX) kinesin-fragment, 2) PEX-FKBP (FK506 binding protein), and 3) a myoV motor fused to an FRB (FKBP-rapamycin-binding) domain (Fig 3A). Addition of a rapamycin-analog (rapalog) induces the hetero-dimerization of FKBP with FRB, and thereby allows for a time-controlled recruitment of the myoV motor to the kinesin-coupled peroxisome. Of note, for this assay we used a myoVb construct (as published in Kapitein *et al*, 2013), which, just like myoVa, showed Ca^2+^-dependent interaction with caldendrin (Fig EV1B, Fig EV3C, Table EV1). Since COS7 cells do not endogenously express caldendrin, we additionally transfected them with an untagged caldendrin construct and confirmed co-expression with the inducible motor system in fixed COS7 cells (Fig EV3D). The inducible dimerization assay revealed the expected inhibition of kinesin-driven peroxisome motility by myoVb recruitment (Fig 3B, Movie EV3.2). The efficiency of myosin-induced anchoring of peroxisomes, as assessed by the mean-squared displacement (MSD) of peroxisomes before and after rapalog addition, was not affected by the simultaneous expression of caldendrin (Fig 3C). We therefore concluded that caldendrin binding to myoV is unlikely to inhibit association of myoV with F-actin.

**Figure 3:**
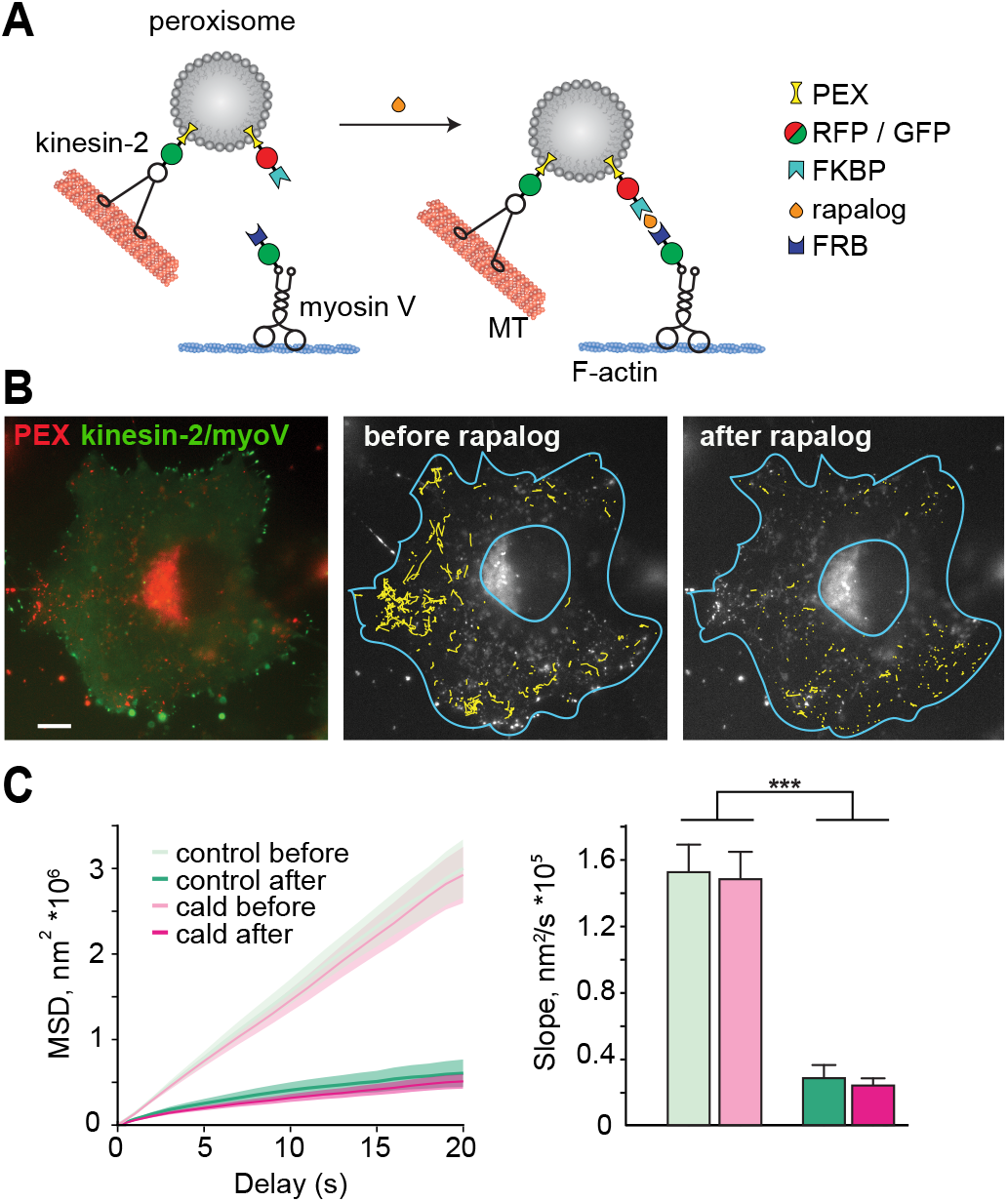
Inducible dimerization assay in COS7 cells shows that the presence of caldendrin does not affect ability of myosin V to bind actin filaments.

**A** Schematic representation of the assay components. Three plasmids are co-expressed in COS7 cells: a constitutively active kinesin-2 motor (KIF17) fused to a GFP-tagged peroxisome targeting sequence (KIF17-GFP-PEX13), an RFP-tagged PEX13 fused to an FKBP protein (PEX-RFP-FKBP), and a constitutively active myosin Vb motor fused to an FRB protein (myoV-GFP-FRB). Addition of the small molecule rapalog leads to dimerization of the FKBP and FRB, thus recruiting myoV to the peroxisome. Additionally, the cells were transfected with untagged caldendrin (or an empty control vector) to assess the effect of caldendrin binding to myoV on its activity in living cells.

**B** Left: Representative image of COS7 cell expressing the inducible dimerization system described in **A**. Red: Peroxisomes (PEX-RFP-FKBP), Green: KIF17-GFP-PEX13 and myoV-GFP-FRB. Scale bar = 10 μm. Middle and right: Individual tracks of kinesin-driven peroxisome movements reconstructed over 20 seconds (yellow) before (middle) and after (right) addition of rapalog indicate that the recruitment of myosin V slows down the motility of peroxisomes. Blue outlines represent manually drawn cellular border and nucleus (excluded from analysis). Also see Movie EV3.1.

**C** Analysis of the mean squared displacement (MSD) of peroxisomes indicates that addition of rapalog induces efficient stalling (2-way ANOVA, *** p<0.0001 for rapalog). However, no difference between control and caldendrin (cald) transfected cells could be observed (2-way ANOVA, n.s. for interaction. Mean ± SEM; n control=14 cells, n cald=23 cells; 3 experiments)

### Loss of caldendrin in hippocampal neurons destabilizes ER inside dendritic spines

Any inhibitory effect of caldendrin on myoV would be especially relevant where caldendrin is present in high concentrations, i.e. in a subset of dendritic spines (Dieterich et al, 2008). At the same time, a process that relies on myoV in hippocampal neurons is the localization of SER, and of the ER-based spine apparatus organelle, to selected spines (Perez-Alvarez *et al*, 2020; Konietzny *et al*, 2019). Spinous targeting of SER can be observed during periods of high synaptic activity, i.e. when Ca^2+^ concentration in an individual synapse is temporarily increased (Perez-Alvarez *et al*, 2020) which also favours activation of caldendrin (Mikhaylova *et al*, 2018). Therefore, we proposed that caldendrin might be involved in the regulation of SER targeting to spines by tuning myoVa activity.

To investigate this, we compared SER-spine-entry dynamics in adult dissociated hippocampal neurons of caldendrin knockout mice to those of their wild-type litter mates. SER-spine-entry during time-lapse imaging (10 min, 0.5 fps) was analyzed by expressing a fluorescent ER marker (ER-tdimer2) together with a cell fill (maxGFP) to visualize the cell outline (Fig 4A). As in this experiment we could not definitively distinguish dendritic spines, harboring a synapse, from filopodia, we collectively refer to them as “protrusions” from here on. The frame-by-frame presence (yes/no) of SER inside a given protrusion was analyzed and protrusions that contained SER during the entire imaging period were scored as having a “stable” SER insertion, others were counted as “transient” or “absent” (Fig 4B). We observed that SER-localization to dendritic protrusions was more dynamic and less uniform in knockout compared to wild type neurons (Fig 4B). In agreement with previous findings, caldendrin knockout neurons tended to have fewer protrusions in general than wild type neurons (Fig 4C; Mikhaylova *et al*, 2018). Further, in caldendrin knockout neurons, the fraction of protrusions that were only transiently visited by the SER was increased (from 22% to 30%, Fig 4D), and transient protrusions had a higher number of SER visits (on average 1.3 in wild type and 2.0 in caldendrin knockout; Fig 4E,F). We interpret these results as an increased number of “failed” anchoring attempts by myoVa in the absence of caldendrin.

**Figure 4:**
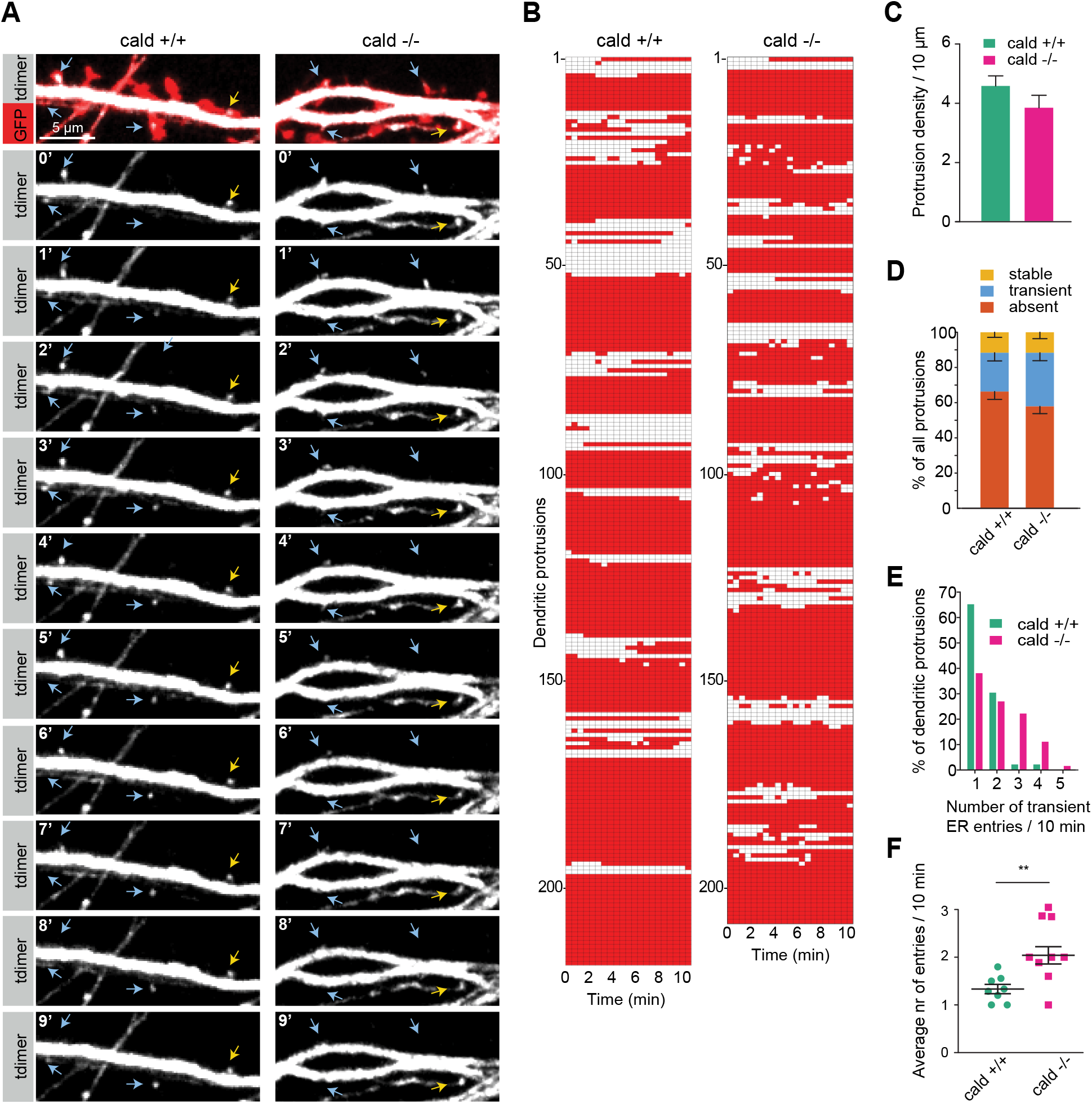
Loss of caldendrin leads to an increased frequency of SER entry and decreased fraction of stable SER in dendritic protrusions of mouse primary hippocampal neurons.

**A**10 min time-lapse imaging of DIV12 wild-type (cald +/+) or caldendrin knock-out (cald −/−) mouse primary hippocampal neurons expressing a cell fill (maxGFP) and an ER-marker (ER-tDimer). Yellow arrows indicate protrusions with a stably anchored ER (>10 min). Blue arrows indicate protrusions that show transient presence of ER. Scale bar = 5 μm.

**B** Visualization of SER presence over time in individual protrusions of wild-type (cald +/+) or caldendrin knock-out (cald −/−) neurons as shown in **A**. Red = no SER present, white = SER present.

**C** Quantification of protrusions density in cald +/+ and cald −/−neurons shows a non-significant reduction in the knock out neurons. Data shown as mean ± SEM.

**D – E** Quantification of dynamic SER presence in protrusions of cald +/+ and cald −/−neurons.

**D** Protrusions of cald −/−neurons show a non-significant trend towards more protrusions with transiently present ER at the expense of protrusions where the SER was absent throughout the imaging period. (mean ± SEM, n.s. 2-way ANOVA, n wt = 8, 1 experiment; n KO = 9; 1 experiment).

**E** In protrusions with a transiently present SER, cald −/−neurons have an increased number of ER visits over 10 minutes compared to the wild-type.

**F** Quantification of the data in **E** across individual cells shows a significant higher number of SER visits in cald −/−neuron protrusions (mean ± SEM, ** p = 0.0078, unpaired t-test, n wt = 8, n KO = 9; 1 experiment).

### Overexpression of caldendrin in hippocampal neurons leads to an increase of SER localized inside dendritic spines

In a complementary set of experiments, we over-expressed GFP-tagged caldendrin in dissociated rat hippocampal neurons together with ER-tdimer2 for 18-24 hours to observe SER-spine-entry dynamics in the presence of excess caldendrin (Fig 5A). Again conducting live time-lapse imaging (10 min, 0.5 fps), we found that this time, the distribution of “stable”, “transient” and “absent” protrusions had significantly changed, with an increase in “stable” protrusions from 9 % to 17 % (Fig 5B, Fig EV5A,B, Movie EV5). In order to distinguish between spine-localized and filopodia-localized SER, we used immunostaining on fixed cells to visualize ER together with the post-synaptic marker homer1 (Fig 5C). Complementary to the caldendrin knockout, caldendrin overexpression caused an increase in both spine and filopodia density (Fig 5D,E). Importantly, caldendrin overexpression also significantly increased the percentage of homer1-positive spines that contained SER (19.9% to 24.5%, Fig 5F).

**Figure 5:**
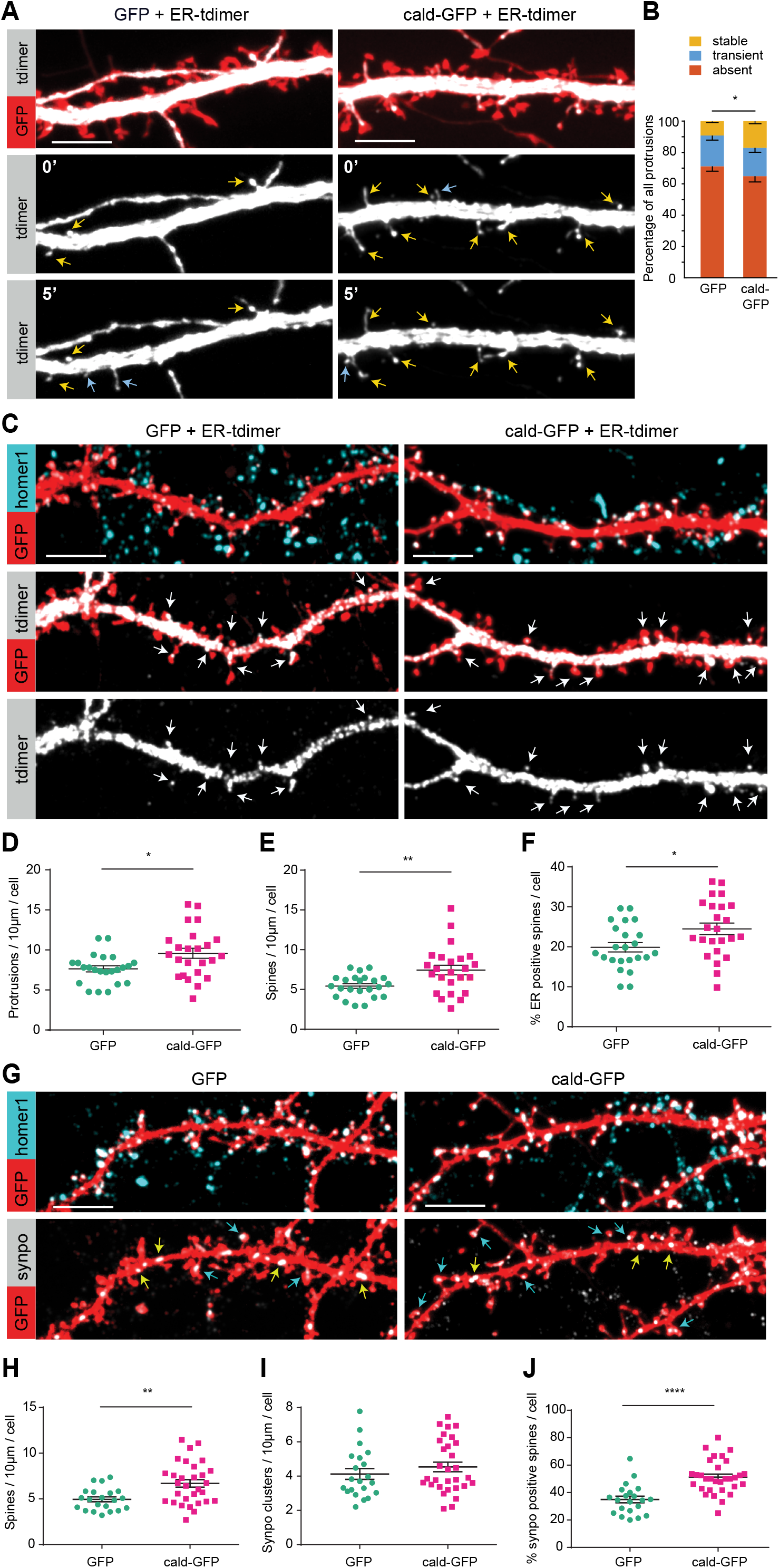
Overexpression of caldendrin leads to an increase of stable SER in dendritic spines of rat primary hippocampal neurons.

**A** Example confocal images from 10 min time-lapse imaging of control (GFP) or caldendrin (cald-GFP) transfected rat primary hippocampal neurons (DIV17) expressing an ER-marker (ER-tDimer). Yellow arrows show dendritic protrusions with a stably inserted ER (>10 min). Blue arrows indicate protrusions with a transient ER entry. Scale bar = 5 μm. Also see Movie EV5.

**B** Quantification of dynamic SER presence in protrusions of control (GFP) or caldendrin (cald-GFP) transfected neurons. In protrusions of cald-GFP neurons, the percentage of stably localized ER (> 10 min) is increased, while transient SER visits are decreased compared to the control. (mean ± SEM, 2-way ANOVA, * p=0.0386 for interaction, n control=9, n cald=10; 2 experiments)

**C** Representative confocal images of control (GFP) or caldendrin (cald-GFP) transfected rat primary hippocampal neurons expressing an ER-marker (ER-tDimer). To pre-load overexpressed caldendrin with Ca^2+^, neurons were stimulated with 50 μM bicuculline for 5 min 8 hours prior to fixation and stained with a homer1-antibody to visualize synapse-containing spines. Arrows indicate spines that contain ER. Scale bar = 5 μm.

**D – F** Quantification of protrusion- and spine density and the number of SER-positive spines as shown in **C**. (mean ± SEM, n GFP ctrl = 24 cells, n Cald-GFP = 25 cells; 3 experiments).

**D** The density of dendritic protrusions (spines and filopodia) is increased in cald-GFP overexpressing neurons compared to the control. (* p=0.0119, unpaired t-test)

**E** The density of homer1-positive dendritic spines is increased in cald-GFP overexpressing neurons compared to the control. (** p=0.0049, unpaired t-test)

**F** The percentage of ER-containing dendritic spines is increased in cald-GFP overexpressing neurons compared to the control. (* p=0.0179, unpaired t-test)

**G** Representative confocal images of control (GFP) or caldendrin (cald-GFP) transfected rat primary hippocampal neurons pre-treated with bicuculline as in **C** and stained against homer1 and the spine apparatus-marker synaptopodin. Yellow arrows indicate synaptopodin clusters localized inside the dendritic shaft, blue arrows indicate spine-localized synaptopodin. Scale bar = 5 μm.

**H – J** Quantification of spine density, synaptopodin clusters and synaptopodin-positive spines as shown in **G**. (mean ± SEM, n GFP ctrl = 21 cells, n cald-GFP = 30 cells; 2 experiments).

**H** Spine density is increased in cald-GFP overexpressing neurons compared to the control (in line with 5e). (** p = 0.0024, unpaired t-test)

**I** The total number of synaptopodin clusters is not affected by caldendrin overexpression. (n.s., unpaired t-test)

**J** The percentage of synapotpodin-positive spines is increased in cald-GFP overexpressing neurons compared to the control. (**** p < 0.0001, unpaired t-test).

A recent study reported that about 90 % of spines that contain stably inserted ER over a time period of 5 hours or longer are also positive for the spine apparatus marker protein synaptopodin (Perez-Alvarez *et al*, 2020). Additionally, we have previously shown that enrichment of synaptopodin, and the spine apparatus formation, depend on the activity of myoVa (Konietzny *et al*, 2019). We therefore hypothesized that overexpression of caldendrin would affect the localization of synaptopodin and the spine apparatus at dendritic spines.

To show this, we expressed caldendrin-GFP (or GFP control) in rat hippocampal neurons and stimulated with 50 μM bicuculline for 15 min 8 hours prior to fixation to increase synaptic activity, as induction of long-term potentiation (LTP) of synapses has been shown to boost spine-apparatus formation (Chirillo *et al*, 2019). To visualise synapses and the spine apparatus, neurons were stained for homer1 and synaptopodin, respectively (Fig 5G). Once again, caldendrin-GFP overexpression increased spine density (Fig 5G, H), while it did not affect the overall density of synaptopodin clusters throughout the dendrite (Fig 5I). Interestingly, we found that the percentage of synaptopodin-positive spines had increased from 34.9 % in GFP control, to 51.3 % in caldendrin-GFP.

These results are in line with an increase of stably inserted SER observed in the live imaging experiments (Fig 5B). From this we conclude from this that overexpression of caldendrin increases the percentage of spines that contain a spine apparatus. Altogether, these results suggest that caldendrin plays a role in stabilizing the presence of SER in the spine head by stabilizing myoV in its anchoring function.

## Discussion

In this study, we aimed to decipher how the modalities of Ca^2+^-sensing in myoV influence the processivity and stalling behaviour of the motor protein, which is instrumental for the proper synaptic targeting of SER to a subset of active dendritic spines in hippocampal neurons.

Although it is well accepted that Ca^2+^/CaM regulates the active vs. inactive conformation and processivity of myoV and VI, there are surprisingly few studies addressing the exact role of Ca^2+^ in regulation of unconventional myosins in cells (Krementsov *et al*, 2004; Wang *et al*, 2008). Here, we found through a mass spectrometry screen that another brain-specific Ca^2+^-binding protein, caldendrin, associates with different myosin family members, including myoVa, Vb and myoVI, in a Ca^2+^-dependent manner. The moderate Ca^2+^-binding affinity of the protein (Reddy et al., 2014) would restrict Ca^2+^-dependent interactions of caldendrin to compartments with steep Ca^2+^-gradients, such as dendritic spines. Additionally, in contrast to the diffusely distributed CaM, caldendrin is enriched in big and mature spines that are expected to experience frequent synaptic inputs. Known roles of caldendrin include activity-dependent stabilization of F-actin via interaction with cortactin, regulation of IP_3_Rs present at the ER, as well as regulation of L-type calcium channels and synapse-to-nucleus communication via the messenger-protein Jacob (Dieterich *et al*, 2008; Li *et al*, 2013; Tippens & Lee, 2007; Mikhaylova *et al*, 2018). Larger mushroom-like spines are stable over time, have a stronger synaptic connection, and frequently contain SER. MyoV has been shown to be implicated in the activity-dependent targeting of SER upon induction of synaptic plasticity (Kneussel & Wagner, 2013; Perez-Alvarez *et al*, 2020). We therefore decided to focus on myoV and asked whether the interaction between caldendrin and myoV isoforms could have functional consequences for targeting and stabilization of the ER-derived structures in dendritic spines.

To first identify the binding interface, we performed co-immunoprecipitation assays with various truncation constructs of both proteins. Ca^2+^-dependent accessibility of caldendrin is mediated by its C-terminal domain, which masks the N-terminus in the absence of Ca^2+^. However, we still observed a Ca^2+^-dependent interaction between the isolated caldendrin N-terminus and myoV, which suggested that another calcium sensing modality might be involved. By further narrowing down the binding region, we found that caldendrin specifically recognizes a 49 amino-acid fragment of myoV containing part of the lever arm and the first IQ-motif, IQ1. The fact that the caldendrin and IQ1-CaM binding sites are either in close proximity or possibly even overlap carries interesting implications for the effect of caldendrin binding on myoV activity. IQ1-CaM was shown to be responsible for the observed stimulation of actin-activated ATPase activity of myoV by micromolar Ca^2+^: in its inactive state, myoV is in an auto-inhibited, folded conformation in which the C-terminal cargo binding domain is in contact with the N-terminal motor domain (Sellers *et al*, 2008; Wang *et al*, 2008). Ca^2+^-induced conformational change of IQ1-CaM can break the interaction between motor and cargo domain, thus activating motor function (Lu *et al*, 2012; Shen *et al*, 2016). A possible explanation for the observed Ca^2+^-sensitivity of the caldendrin-myoV interaction could therefore be the described conformational change in IQ1-CaM upon binding of Ca^2+^ (Lu *et al*, 2012; Shen *et al*, 2016).

In addition to that, Ca^2+^ binding to IQ1-CaM was shown to not only cause significant changes to the lever arm, but also propagated changes that extend into the motor domain (Trybus *et al*, 2007), which might explain the observed reduction of myoV velocity between Ca^2+^/CaM and apoCaM (Krementsov *et al*, 2004). It is therefore likely that caldendrin binding closely to the IQ1-CaM interface will affect myoV motor function. Indeed, *in vitro* competition pull downs with CaM and caldendrin indicated that, in the presence of Ca^2+^, caldendrin can replace CaM at IQ1. Importantly, full decoration of the lever-arm with CaM is a known prerequisite for processive myoV motor function, as detachment of one or several CaMs leads to destabilization of the lever arm, which is then no longer able to translate the force generated by ATP hydrolysis into processive motility (Koide *et al*, 2006; Trybus *et al*, 2007). It is possible that potential displacement of IQ1-CaM by caldendrin would affect myoVa processivity in a similar way. Here, we further showed that processive myoVa motor activity is generally inhibited by the presence of caldendrin in a reconstituted *in vitro* gliding assay. The exact mechanism of this inhibition remains to be elucidated. However, it is known that in the gliding assay, myosin motors that are attached to the glass surface are forced into an “open” conformation, preventing formation of the auto-inhibited state (Krementsov *et al*, 2004). We therefore conclude that the inhibitory effect that caldendrin has on myoVa affects the motor in its open and active conformation.

As conformational changes of CaM and/or the myoV lever arm can extend to the motor domain (Trybus *et al*, 2007), we reasoned that the observed inhibition of myoVa processivity in the gliding assay could be due to caldendrin affecting myoVa interaction with F-actin. However, using an *in vitro* sedimentation assay and an inducible motor-cargo assay in COS7 cells, we did not find any evidence that caldendrin negatively affects myoV binding to F-actin. Alternatively, caldendrin could exert its inhibitory function by “locking” the lever arm in place, preventing myoV from stepping forward, but without displacing it from F-actin. This would explain both results from the *in vitro* assay and the inducible motor-cargo assay.

The invasion of SER into activated dendritic spines is one example of a cellular process where controlled stop-and-go behaviour of myoV plays an instrumental role. In a recent study from the laboratory of T.G. Oertner it has been shown that transient SER spine entries are not random but target highly activated spines (Perez-Alvarez *et al*, 2020). In another study, LTP-driven transformation of the SER into a spine apparatus has been reported (Chirillo *et al*, 2019). Accordingly, in our previous work we have shown that myoV activity is required for synaptic accumulation of synaptopodin and formation of the spine apparatus. Based on this, we hypothesize that the Ca^2+^-dependent interplay of CaM, caldendrin and myoV represents the core mechanism of recruitment and stabilisation of the ER in dendritic spines.

While performing live imaging of SER-spine-entry, we found that dendritic protrusions of caldendrin knockout neurons experienced a higher number of transient SER visits, while the percentage of stably inserted SER was similar to the wyld type control. This could be explained by myoV being “over-active” in the absence of its inhibitor caldendrin. In this scenario, SER protrusions would be repeatedly targeted to spines by an over-active myoV, but then fail to become stably anchored. Interestingly, in a complementary experiment using caldendrin overexpression, we observed an increase of stable SER, while the dynamics of transient SER visits were unchanged compared to the control. We therefore propose a model in which caldendrin stops the processivity of myoV and arrests it in place together with its cargo. Ca^2+^ concentrations are highest in the heads of large dendritic spines that usually experienced high frequency synaptic stimulation, for instance during LTP. As caldendrin has to compete with other Ca^2+^-binding proteins for Ca^2+^ ions, it is most likely to become activated in the confined spine head, and to a lesser extent in the spine neck or dendritic shaft (Raghuram *et al*, 2012; Reddy *et al*, 2014). We speculate that, as myoV processively transports SER into activated spines, once it reaches the spine head Ca^2+^/caldendrin will replace Ca^2+^/CaM from IQ1 and inhibit further motility. Importantly, it does not dissociate myoV from F-actin, but rather locks it in place to function as a tether.

Further, following synaptic activation, we observed an increased number of dendritic spines containing SER as well as the spine apparatus-marker synaptopodin, compared to mock treated cells. According to our model, we suggest that caldendrin overexpression led to an increased dwell-time of SER inside dendritic spines, which through association with synaptopodin could then progressively trigger spine apparatus formation (Konietzny *et al*, 2019).

What is the role of SER inside dendritic spines? In recent work it has been proposed that the presence of SER correlates with LTP-like activity of individual spines, and that it is involved in the depotentiation of spines in an mGluR1-dependent manner (Perez-Alvarez *et al*, 2020; Holbro *et al*, 2009). SER contains IP_3_Rs which are important for the long-term depression (LTD) and downscaling of over-activated spines (Holbro *et al*, 2009; Nakamura *et al*, 1999; Segal & Korkotian, 2014). Interestingly, caldendrin binds IP_3_R and decreases its sensitivity for IP_3_ (Li *et al*, 2013). This could provide a mechanism to tune the sensitivity of caldendrin- and SER-positive spines to LTP- and LTD-like stimuli. Indeed, in our previous work we found that caldendrin knockout mice fail to form a stable LTP in CA1-Schaffer collaterals. This effect could be partially rescued by stabilization of actin filaments with jasplakinolide during the LTP induction (Mikhaylova *et al*, 2018). It is possible that the deficiency in SER anchoring via myoV was another limiting factor for the maintenance of synaptic potentiation. Altogether, we propose a Ca^2+^-sensitive module comprising myoV, CaM and caldendrin as a key cargo-delivery mechanism that is operational in a subset of dendritic spines and contributes to the diversification and specification of excitatory glutamatergic synapses.

## Materials and Methods

### Animals

Wistar Unilever HsdCpb:WU (Envigo) rats and caldendrin knockout mice (Mikhaylova *et al*, 2018) were used in this study. All animal experiments were carried out in accordance with the European Communities Council Directive (2010/63/EU) and the Animal Welfare Law of the Federal Republic of Germany (Tierschutzgesetz der Bundesrepublik Deutschland, TierSchG) approved by the local authorities of Sachsen-Anhalt/Germany (reference number 42502-2-987IfN and 42502-2-1264 LIN, TV 42502-2-1009 UniMD) or of the city-state Hamburg (Behörde für Gesundheit und Verbraucherschutz, Fachbereich Veterinärwesen) and the animal care committee of the University Medical Center Hamburg-Eppendorf.

### Primary neuronal culture and transfections

Primary hippocampal rat cultures were prepared essentially as described previously (Kapitein *et al*, 2010). In brief, hippocampi were dissected from E18 embryos, treated with trypsin (0.25 %, *Thermo Fisher Scientific*) for 10 min at 37°C, physically dissociated by pipetting through a syringe, and plated on poly-L-lysine (*Sigma-Aldrich*, #P2636) coated glass coverslips (18 mm) at a density of 40000-60000 cells per 1 ml in DMEM (*Gibco*, #41966-029) supplemented with 10 % fetal calf serum (*Gibco*, #10270) and antibiotics (*Thermo Fisher Scientific*, #15140122). After 1 h, the plating medium was replaced by BrainPhys neuronal medium supplemented with SM1 (*Stem Cell kit*, #5792) and 0.5 mM glutamine (*Thermo Fisher Scientific*, #25030024). Cells were grown in an incubator at 37 °C, 5 % CO_2_ and 95 % humidity.

Cultures of mouse primary hippocampal neurons were prepared as described previously (Mikhaylova *et al*, 2018). In brief, hippocampi were dissected from male and female P0 mouse pups and after 10 min treatment with trypsin at 37 °C cells were physically dissociated by pipetting through a syringe. Neurons were plated on glass coverslips (18 mm) coated with poly-L-lysine (*Sigma-Aldrich*) at a density of 30.000 cells per ml in DMEM (*Gibco*) supplemented with 8 % FCS and 1% penicillin/streptomycin. Following attachment, mouse neuronal cultures were kept in Neurobasal medium (NB; *Gibco*) supplemented with 2 mM glutamine, 1 % penicillin/streptomycin and 1X B27 supplement (*Gibco*), at 37 °C, 5 % CO2 and 95 % humidity. Primary cultures were transfected with lipofectamine 2000 (*Thermo Fisher Scientific*) according to the manufacturer’s instructions. For co-transfection of plasmids the ratios of different constructs were optimized individually, and optionally by addition of an “empty vector” (pcDNA3.1), to tune expression levels. Before transfection, the conditioned neuronal medium was removed from the cells and kept at 37°C. Neurons were transfected in BrainPhys medium lacking SM1 (rat cultures) or B27 (mouse cultures) by incubation in the transfection mixture for 45 min – 1.5 h. After transfection, the medium was exchanged back to the conditioned medium. Experiments on transfected neurons were performed 24 hours after transfection.

For fixed imaging of ER spine localization together with synaptopodin, 16 hours after transfection and 8 hours before fixation, neurons were transferred into medium containing 50 μM Bicuculline (*Tocris*) for 15 min to increase synaptic activity, and then back to their conditioned medium.

### Culturing and transfection of cell lines

HEK293T cells (DSMZ, ACC 635) and COS7 cells (ATCC, CRL-1651) were maintained in full medium consisting of Dulbecco’s modified Eagle’s medium (DMEM; GIBCO, Thermo Fisher) supplemented with 10 % fetal calf serum (FCS), 1 × penicillin/streptomycin and 2 mM glutamine at 37 °C, 5 % CO2 and 95 % humidity. For the expression of biotinylated proteins, cells were grown in full medium made with a 50 % DMEM, 50 % Ham’s F-10 Nutrient Mix (GIBCO, Thermo Fisher) mixture. HEK293T cell transfections were done using MaxPEI 25K (*Polysciences*) in a 3:1 MaxPEI:DNA ratio according to the manufacturer’s instructions. Transfected HEK293T cells were harvested 18-24 hours after transfection. The cells were washed 1 × in cold TBS, resuspended in 2 ml TBS and pelleted for 3 min at 1000g. The cell pellet was lysed in in 500 μl extraction buffer (20 mM Tris pH 8, 150 mM NaCl, 1% Triton-X-100, 5 mM MgCl_2_, complete protease inhibitor cocktail (Roche)), kept on ice for 30 min and centrifuged for 15 min at 14000 × g. The supernatant (cleared lysate) was then further used for experiments.

COS7 cells were plated and imaged in 35 mm glass-bottom μ-dishes (*Ibidi*, #81156) or on uncoated glass coverslips. They were transfected using FuGene HD (*Promega*, #E2311) in a 3:1 FuGene:DNA ratio according to the manufacturer’s instructions. Transfected COS7 cells were imaged 18-24 hours after transfection. During image acquisition, rapalog (A/C Heterodimerizer, *TaKaRa*, #635056) was added at a final concentration of 100 nM.

### Immunoblotting

For immunoblot analysis, samples were either taken up in commercial Bolt LDS sample buffer (Invitrogen), or self-made SDS sample-buffer (4 × stock: 250 mM Tris-HCl, pH 6.8, 8 % (w/v) SDS, 40 % (v/v) glycerol, 5 % (v/v) β-mercaptoethanol, 0.004% bromophenol blue, pH 6.8) as specified. In general, samples were boiled at 98°C for 10-15 min, run on 4 % −20 % acrylamide gradient gels and blotted on PVDF membranes in blotting buffer (192 mM glycine, 0.1 % (w/v) SDS, 15 % (v/v) methanol, 25 mM Tris-base, pH 8.3). After blocking the membranes in 5 % skim milk in Tris-buffered saline (TBS, 20 mM Tris pH 7.4, 150 mM NaCl, 0.1 % Tween-20), membranes were incubated with primary antibodies diluted in TBS-A (TBS pH 7.4, 0.02 % sodium-azide) overnight at 4°C. Corresponding HRP-conjugated secondary antibodies or HRP-Strepdavidin were applied for 1.5 h at RT in 5 % skim milk in TBS. The membranes were imaged on a ChemoCam imager (*Intas*).

### Biotin-streptavidin pull-down experiments

HEK293T cells were co-transfected with HA-BirA and either Caldendrin-pEGFP-bio, or pEGFP-bio (control vector) and harvested on the next day as described above. Magnetic Streptavidin M-280 Dynabeads (*Invitrogen*) were washed 3 × in washing buffer (20 mM Tris pH 8, 150 mM KCl, 0.1 % Triton-X 100), blocked in 3 % chicken egg albumin (Sigma) for 40 min at RT, and again washed 3 × in washing buffer. The cleared cell lysate was added to the blocked beads and incubated at 4 °C on a rotator overnight. After the incubation period, the beads were washed 2 × with low salt washing buffer (100 mM KCl), 2 × in high salt washing buffer (500 mM KCl) and again 2 × in low salt washing buffer. For preparation of whole mouse brain extract, 9 ml lysis buffer (50mM Tris HCl pH 7.4, 150 mM NaCl, 0.1 % SDS, 0.2 % NP-40, complete protease inhibitor cocktail (*Roche*)) were added per 1 g of tissue weight, and the tissue was lysed using a Dounce homogenizer. The lysate was cleared first for 15 min at 1000 × g, and the supernatant was again centrifuged for 20 min at 15000 g to obtain the final lysate. The washed beads were then combined with 1 ml of the cleared mouse brain lysate and incubated for 1 h at 4 °C on a rotator. The beads were divided into 2 × samples, and incubated with mouse brain lysate that was either substituted with 2 mM EGTA or with 0.5 mM CaCl_2_ and 1 mM MgCl_2_. After the incubation period, the beads were washed 5 × in washing buffer and resuspended in Bolt LDS sample buffer (Invitrogen) for subsequent SDS-PAGE and mass-spectrometry or western blot analysis. For mass-spectrometric analysis, the samples were separated on a commercial Bolt Bis-Tris Plus Gel (*Invitrogen*) and the intact gel was sent to Erasmus MC Proteomics Center, Rotterdam, for mass spectrometry analysis (see below).

### Co-immunoprecipitation from HEK293T cells

HEK293T cells were transfected with the respective GFP-tagged myosin construct and either with tagRFP-tagged caldendrin or tagRFP as a control, and harvested as described above. 45 μl of the cleared lysate were taken as an “input” sample. The remaining SN was split into 2 × 200 μl, distributed into separate tubes and substituted with either 2 mM EGTA or 0.5 mM CaCl_2_ and 1 mM MgCl_2_. GFP-trap beads (*Chromotek*) were washed 2 × in extraction buffer and an equivalent of 7 μl slurry was added to the tubes containing the cell lysates. The beads were then incubated on a rotor at 4°C for 2 – 4 hours. After the incubation period, the beads were washed 3 × in washing buffer (20 mM Tris, 150 mM NaCl, 0.5 % Triton-X-100, complete protease inhibitor cocktail (*Roche*)) that was either substituted with 2 mM EGTA or 0.5 mM CaCl_2_ and 1 mM MgCl_2_. The beads were then taken up in 30 μl SDS sample-buffer for SDS-PAGE.

### Pull down of purified myosin with recombinant caldendrin

HEK293T cells were transfected with the respective GFP-tagged myosin constructs and either with tagRFP-tagged caldendrin or tagRFP as a control, and harvested as described above. The cleared lysate was incubated with an equivalent of 7 μl GFP-trap slurry (*Chromotek*) for 6 h at 4°C on a rotor, then the beads were washed 3 × with washing buffer (20 mM Tris, 150 mM NaCl, 0.5 % Triton-X-100, complete protease inhibitor cocktail (*Roche*)), blocked with washing buffer supplemented with 5 % BSA, and then divided into 4 equal samples. Each one was incubated with washing buffer supplemented with 100 nM of either purified, recombinant wild type caldendrin, or a caldendrin calcium binding mutant (D243A, D280A, (Mikhaylova *et al*, 2018)), in the presence or absence of 2 mM EGTA or 100 μM CaCl2. After the incubation period, the beads were washed 3 × in washing buffer (20 mM Tris, 150 mM NaCl, 0.5 % Triton-X-100, complete protease inhibitor cocktail (*Roche*)) that was either substituted with 2 mM EGTA or 100 μM CaCl2. The beads were then taken up in 30 μl SDS sample-buffer for SDS-PAGE.

### F-actin co-sedimentation assay

This protocol was adapted from *Cytoskeleton #BK037* assay kit. HEK293T cells were co-transfected with GFP-tagged full-length myosin Va and either with tagRFP-tagged caldendrin or tagRFP as a control. After 24 hours, the cells were mechanically lysed in room-temperature F-actin stabilization buffer (20 mM Tris pH 7.4, 20 μM CaCl2, 100 mM KCl, 2 mM MgCl2, 200 μM ATP, 1 mM DTT, 200 nM Jasplakinolide (*Tocris*), complete protease inhibitors (*Roche*)) by repeatedly passing through a 25 Gauge syringe. The lysate was incubated at 37 °C for 10 min before centrifuging at 350 g for 5 min to pellet unbroken cells and debris. The supernatant was transferred into fresh tubes and centrifuged at 100.000 g for 1 hour to pellet F-actin. After ultra-centrifugation, pellet and supernatant were separated, solubilized in SDS-sample buffer and analyzed by SDS-PAGE and immunoblotting.

### Mass spectrometry and data analysis

Pull-down samples were separated on a commercial Bolt Bis-Tris Plus Gel (*Invitrogen*) and analyzed at the Erasmus MC Proteomics Center, Rotterdam. Briefly, SDS-PAGE gel lanes were cut into 1-mm slices using an automatic gel slicer. Per sample 8 to 9 slices were combined and subjected to in-gel reduction with dithiothreitol, alkylation with iodoacetamide and digestion with trypsin (*Thermo Fisher Scientific*, TPCK treated), essentially as described by (Wilm *et al*, 1996). Nanoflow LCMS/MS was performed on an EASY-nLC 1000 Liquid Chromatograph (*Thermo Fisher Scientific*) coupled to an Orbitrap Fusion™ Tribrid™ Mass Spectrometer (*Thermo Fisher Scientific*) operating in positive mode and equipped with a nanospray source. Peptide mixtures were trapped on a nanoACQUITY UPLC C18 column (100Å, 5 μm, 180 μm × 20 mm, Waters). Peptide separation was performed on a ReproSil C18 reversed phase column (*Dr Maisch GmbH*; 20 cm × 100 μm, packed in-house) using a linear gradient from 0 to 80% B (A = 0.1 % formic acid; B = 80% (v/v) acetonitrile, 0.1 % formic acid) in 60 min and at a constant flow rate of 200 nl/min. Mass spectra were acquired in continuum mode; fragmentation of the peptides was performed in data-dependent mode.

Mass spectrometric raw data were analyzed using the MaxQuant software suite (version 1.5.4.1) for identification and relative quantification of peptides and proteins. A false discovery rate (FDR) of 0.01 for proteins and peptides, and a minimum peptide length of 6 amino acids were required. The Andromeda search engine was used to search the MS/MS spectra against the *Homo sapiens* and *Mus musculus* Uniprot databases (release July 2016) concatenated with the reversed versions of all sequences and a contaminant database listing typical background proteins. A maximum of two missed cleavages were allowed. MS/MS spectra were analyzed using MaxQuant’s default settings for Orbitrap and ion trap spectra. The maximum precursor ion charge state used for searching was 7 and the enzyme specificity was set to trypsin. Further modifications were Cysteine carbamidomethylation (fixed) as well as Methionine oxidation (variable). The minimum number of peptides for positive protein identification was set to 2. The minimum number of razor and unique peptides was set to 1. Only unique and razor non-modified, methionine oxidized and protein N-terminal acetylated peptides were used for protein quantitation.

### Analysis of mass spectrometry data

Identified peptides (see Table 1) were manually curated by removing ambiguous peptides if more than one of their associated proteins also had other unique peptides. Sample variability of the amount of “bait” (caldendrin-GFP or GFP alone) was taken into account by summing the intensities of the highest (>60th percentile) intensity GFP-peptides in each sample and using this as a relative correction factor for all other peptide intensities in that sample. Intensity ratios of proteins with at least n=5 associated peptides in this curated list were then calculated according to 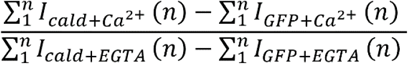.

All protein intensity data were manually verified to accurately reflect the calculated ratio.

### Constructs and cloning

All constructs were verified by sequencing. A complete list of expression constructs used in this study is provided in the Appendix.

The TwinStrep-GFP-myosin constructs were produced by amplifying the TwinStrep-tag sequence by PCR from a TwinStrep-mCherry empty vector template (gift from A. Aher, Utrecht University), and ligating it into the pmEmerald-myoV vector using the NheI cutting site. GFP-myosin fragments for co-immunoprecipitations were amplified from the pmEmerald-myoV template by PCR. The whole myoV sequence was cut from the vector using the NotI cutting site, and the PCR amplified fragments were re-inserted into the vector using sequence overlap and the Cold Fusion Cloning Kit (*SBI, MC010B-1*). The minimal binding region between caldendrin and myoV was narrowed down to a 49 amino-acid fragment containing the first <u>IQ motif</u> (aa742-791, FGKTKIFFRAGQVAYLEKLRAD<u>KLRAACIRIQKTIRGWLLRK RYL</u>CMQR).

The caldendrin-tagRFP constructs were made by amplifying full length or caldendrin fragments from the pcDNA3.1/caldendrin vector and pasting them into the tagRFP-N plasmid *(Evrogen, #FP142*) using EcoRI and BamHI restriction and ligation.

pET/Calmodulin construct: Untagged human calmodulin was amplified from *Addgene* plasmid #47603 via PCR and cloned into an empty pET/T7 expression vector (gift from H.J. Kreienkamp, UKE Hamburg) using NdeI and HindIII restriction and ligation.

### Immunocytochemistry

Cells were fixed in 4 % Roti-Histofix (*Carl Roth*), 4 % sucrose in PBS for 10 min at RT and washed three times with PBS, before they were permeabilized in 0.2 % Triton X-100 in PBS for 10 min. The cells were then washed 3 × in PBS and blocked for 45 min at RT with blocking buffer (BB / 10 % horse serum, 0.1 % Triton X-100 in PBS). Incubation with primary antibodies was performed in BB at 4 °C overnight. After 3 × washing in PBS, cells were incubated with corresponding secondary antibodies in BB for 1 h at RT. Finally, coverslips were washed 3-5 × 10 min in PBS and mounted on microscope slides with Mowiol. Mowiol was prepared according to the manufacturer’s protocol (9.6 g mowiol 4-88 (*Carl-Roth*), 24.0 g glycerine, 24 ml H_2_O, 48 ml 0.2 M Tris pH 8.5, including 2.5 g Dabco, (*Sigma-Aldrich* D27802).

### Fixed and live cell imaging: confocal microscopy

Z-stack images of fixed primary hippocampal neurons were acquired either on Leica TCS SP8 and Leica TCS SP5 confocal microscopes using a 63.0 × 1.40 oil objective, or on a Nikon Eclipse Ti-E spinning-disc confocal microscope controlled by VisiView software using a Nikon 100 × 1.49 oil objective using 488 nm, 568 nm and 633 nm excitation lasers. The pixel size was set to 85-90 nm and z-steps varied between 250-350 nm. For the shown representative confocal images, a Gaussian filter (radius 0.5 px) was applied in ImageJ to reduce the visible background noise.

Live imaging of mouse neurons was performed on Leica SP5 and SP8 microscopes, at 5% CO_2_ and 37°C. Imaging settings were as follows: 63x objective (HCX PL APO CS 63.0×1.40 oil objective) and 3x digital zoom, 160.49×160.49 nm size, bidirectional scan, time interval 30 sec, imaging for 10 min, Z-stack with 1 μm step size. Maximum projections of the Z stacks were used for analysis.

### Live cell imaging: wide field and spinning disc microscopy

Wide-field and spinning-disc confocal microscopy was performed with a Nikon Eclipse Ti-E controlled by VisiView software. Samples were kept in focus with the built-in Nikon perfect focus system. The system was equipped with a 60x (*Nikon*, *P-Apo* DM *60x/1.40 oil)* and a 100x TIRF objective (*Nikon*, ApoTIRF 100x/1.49 oil), and 488 nm, 561 nm, and 639 nm excitation lasers. Lasers were coupled to a CSU-X1 spinning disk unit via a single-mode fiber. Emission was collected through a quad band filter (*Chroma*, ZET 405/488/561/647m) followed by a motorized filter wheel (*Prior Scientific*) with filters for CFP (480/40m), GFP (525/50m), YFP (535/30m), RFP (609/54m) and Cy5 (700/75m, *Chroma*) and captured on an Orca flash 4.0LT CMOS camera (*Hamamatsu*). COS7 cells were imaged with the 60x objective, neurons were imaged with the 100x objective. Time-lapse images were acquired sequentially with specified intervals.

### Inducible dimerization assay in COS7 cells

COS7 cells were transfected with four plasmid constructs: KIF17(1-547)-GFP-PEX26, PEX-mRFP-FKBP, MyoVb(1-1090)-GFP-FRB, and either untagged caldendrin, or an empty control plasmid. Imaging was done 18-24 hours after transfection in regular growth medium. If cells were grown on coverslips, they were placed in an attofluor cell chamber (*Thermo Fisher Scientific*). Correct temperature (37 °C), CO_2_ (5 %) and humidity (90 %) were maintained with a top stage incubator and an additional objective heater (*Okolab*). Cells with sufficient expression levels (judged by peroxisome visibility (RFP), peroxisome motility (KIF17) and the presence of myosinVb-accumulation in the cell periphery (GFP)) were selected for time-lapse live imaging. Selected cells were imaged for 20 seconds with a high framerate (1 fps) before any treatment to establish baseline peroxisome motility. Then, Rapalog (A/C Heterodimerizer, *TaKaRa*, #635056) was added manually to a concentration of 100 nM to the cell medium. After a 10 min incubation, cells were imaged again for 20 seconds with 1 fps to determine the effect of the rapalog treatment.

### Analysis of peroxisomal motility

Single particle tracking was performed with TrackMate (Tinevez *et al*, 2017) in Fiji (ImageJ 1.52p) with the following parameters: 300nm particle diameter, 25-to 50-fold threshold, median filter, 1500nm gap, max. 2 empty, 3000nm jump. In MATLAB, both the cell outline and nuclear exclusion region were manually drawn. Only tracks whose average positions were within the cell outline and outside of the exclusion region were included in subsequent mean square displacement (MSD) calculations (msdanalyzer, https://github.com/tinevez/msdanalyzer). The weighted mean of all particle MSDs was calculated for each cell. Cells were only included in the final dataset if Rapalog decreased particle displacement at 20s delay by at least 50%. Exclusion of cells was similar between control and caldendrin expressing cells. A linear fit to each weighted mean was calculated to obtain the slope.

### Protein expression and purification

#### Calmodulin purification from E. coli

A single colony of *E.coli* BL21 transformed with the pET/CaM construct was used to inoculate a 5-ml LB pre-culture. After overnight growth at 30 °C, 2 ml of the pre-culture was used to inoculate a 1-l culture in LB medium and grown at 30 °C for 4 h until the absorbance reached 0.5 at 600 nm. Expression of the plasmid was induced by adding 500 μM IPTG, the culture was put at 16°C and grown for about 16 h. Cells were then harvested at 5000 g for 5 min. The pellet was resuspended in 40 ml lysis buffer (25 mM Tris, 5 mM CaCl2, 2 mM MgCl2, 200 μg/ml lysozyme, 2 μg/ml DNAse) and put on a rotor for 10 min at RT. The lysate was then put on ice and sonicated for 2 × 3 min with a 7 mm sonicator at 30 % with cycle set to 3 × 10 %. After centrifugation at 40.000 g for 35 min, the lysate was passed over a 5 ml HiTrap Phenyl FF (LS) column using the *ÅKTA Start* chromatography system. Flow rate was set to 3 ml/min and pressure to 0.03 mPa. Bound calmodulin was eluted with 25 mM Tris, 10 mM EDTA buffer. Binding and elution was performed 3 x. In between, the column was washed with 25 mM Tris, 5 mM CaCl2 buffer. The eluate was captured in 4 ml fractions, with the peak appearing in fraction 2. Fraction 2 from all 3 elution steps was pooled and dialysed O/N at 4°C against 2 l of 25 mM Tris, 50 mM NaCl, 2 mM CaCl2 buffer. After the dialysis, the eluate was up-concentrated to 3 ml using an *Amicon Ultra* 3000 MW centrifugal filter unit (*Merck*). Concentration assessment with a NanoDrop (∊ = 2980, kDa = 17) resulted in an estimated concentration of 15 mg/ml.

#### Caldendrin purification from E.coli

A single colony of *E.coli* BL21 transformed with the pMXB10/Caldendrin-Intein-chitin binding domain-tagged construct was used to inoculate a 20 ml LB pre-culture. After overnight growth at 37 °C, the whole pre-culture was used to inoculate a 1 l culture in LB medium and grown at 37 °C for 2 h until the absorbance reached 0.5 at 600 nm. Expression of the plasmid was induced by adding 400 μM IPTG, the culture was put at 18 °C and grown O/N for about 16 h. Cells were then harvested at 5000 g for 5 min. The pellet was resuspended in 40 ml intein buffer (20 mM Tris-HCl pH 8.5, 500 mM NaCl, 1 mM EDTA, 0.5 mM TCEP (tris(2-carboxyethyl)phosphine), 1 × complete protease inhibitors). Then approx. 20 mg lysozyme (100.000 U/mg) was added and after 30 min incubation on ice the lysate was sonicated for 3 × 10 sec with a 5 mm sonicator at 30 % and cycle set to 80 %. After the sonication, 50 U/ml benzonase (Santa Cruz) was added, the lysate was incubated on ice for 1 h and then centrifuged at 20.000 g for 25 min. The supernatant containing the protein was incubated with 5 ml chitin resin on a rotor at 4 °C for 2 h. The protein-bound resin was then put in a column, washed with 100 ml intein buffer and with 15 ml elution buffer (intein buffer supplemented with 50 mM DTT), before incubation with 7 ml elution buffer O/N at 4 °C. The eluate was collected approx. 16 h later, and dialysed for 2 h against 1 l intein buffer without TCEP, after 2 h the dialysis buffer was exchanged and left again O/N at 4 °C. After dialysis, the eluate was diluted with 8 ml of 100 mM Tris pH 8, up-concentrated in a Amicon Ultra 10.000 MW centrifugal filter unit (Merck) to 2 ml, diluted with 1.1 ml of 100 mM Tris pH 8, again up-concentrated to 1.5 ml, and then ultra-centrifuged at 100.000 g for 15 min. Finally, the eluted protein was aliquoted, flash-frozen in liquid nitrogen and stored at −80°C. Yield ~ 1.5 mg from 1 l culture.

#### Myosin V purification from HEK293 cells

HEK293T cells were transfected with the TwinStrep-pmEmerald/MyosinV construct and harvested as described above. The cell lysate was supplemented with 2 mM EGTA and incubated on a rotor at 4°C with 40 μl magnetic StrepTactin beads (*iba life sciences*) for 1 h. The beads were then washed 2 × with washing buffer (20 mM Tris, 150 mM NaCl, 0.5 % Triton-X-100, 2 mM EGTA, complete protease inhibitor cocktail (*Roche*)), and finally incubated with BXT elution buffer (100 mM Tris ph8, 150 mM NaCl, 1 mM EDTA, 100 mM biotin) for 10 min on ice. After the incubation period, the beads were centrifuged for 2 min at 5.000 g, and the supernatant containing the protein was used for further experiments. Yield ~ 1.5 μg protein from 10 cm HEK cell dish.

### *In vitro* F-actin gliding assay

The setup for the gliding assay was based on (Preciado López *et al*, 2014). Coverslips and microscope slides were cleaned using a “base piranha” solution: Milli-Q water, 30 % hydrogen peroxide and 30 % ammonium hydroxide were mixed in a 5:1:1 volume ratio and heated to 70°C in a glass beaker. Coverslips and microscope slides were placed in the solution using a slide-holder made of Teflon, and incubated for 10 min at 70°C. After the incubation period they were removed and rinsed 5 × with Milli-Q water, and kept in 0.5 M KOH over night or for at least 15 min, before they were dried in an oven at 70 °C. Flow-channels with a width of approx. 4 mm were assembled from the dried microscope slides and coverslips using Para-film as a spacers. The flow channels were coated with PLL-PEG-biotin (*SuSoS AG*), and κ-casein (Sigma) in the following way: 0.1 mg/ml PLL-PEG-biotin in PEM80 buffer (80 mM Pipes pH 6.8, 1 mM EGTA, 4 mM MgCl2) was added into the flow channels and incubated for 1 h at room temperature. The channels were rinsed with 30 μl PEM80 buffer, and 0.5 mg/ml κ-casein in PEM80 was added. After 7-10 min incubation, the channels were rinsed with 50 μl PEM80 and kept in a high humidity chamber until further use.

G-actin (*Tebu bio*) was stored in G-buffer (20 mM Tris-HCl pH 7.4, 20 μM CaCl2, 200 μM ATP, 1 mM DTT) in 25 μM aliquots at −80°C. Aliquots were slowly thawed on ice overnight the day before they were used. Immediately before use, they were diluted 1:1 in G-buffer, centrifuged at 120.000 g for 5 min at 4°C, and the supernatant was used for experiments. To prepare labelled F-actin, G-actin [12.5 μM] was mixed in a ratio of approx. 3.75:1 with alexa-561-labelled actin [10 μM] (*Thermo Fisher*) by mixing 3 μl G-actin with 1 μl alexa-labelled actin. The actin mix was centrifuged at 120.000 g for 5 min at 4°C, the supernatant was taken off and mixed with 6 μl G-buffer to achieve a volume of 10 μl. To this, 10 μl of 2xF-buffer (40 mM Imidazole, 200 mM KCl, 4 mM MgCl2, 400 mM DTT, 12 mg/ml glucose (w/v), 80 μg/ml catalase, 400 μg/ml glucose oxidase, 1.6 mM ATP) was added to achieve a “1x” concentration of the F-buffer. The actin was left to polymerize for 20 min at room temperature in a dark chamber before use.

For the myosin V gliding assay, 2 μl F-actin (as described above) were taken up in 13 μl 1 × F-buffer (20 mM Imidazole, 100 mM KCl, 2 mM MgCl2, 200 mM DTT, 6 mg/ml glucose (w/v), 40 μg/ml catalase, 200 μg/ml glucose oxidase, 0.8 mM ATP) and flown into the flow channel. The filaments were left to settle for 5 min, then 2 μl of freshly purified GFP-myosin V (as above) were taken up in 13 μl 1 × F-buffer and flown into the channel. The channel was left to settle for 5 min, before gliding of actin filaments was imaged. Other components were added to the channel together with myosin in the following final concentrations [stock]: Calmodulin 30 μM [500 μM], Caldendrin 5 μM [75 μM], CaCl_2_ 100 μM [10 mM]. Timelapse imaging was performed on a Spinning-disc confocal with a Nikon Eclipse Ti-E as described under “Live cell imaging”, with 1 frame per second for two to three minutes.

### Analysis of F-actin gliding

In order to assess the percentage of actin filaments that was being actively transported (“gliding”) as opposed to being immobile, the first and 30^th^ frame of a time lapse stack were overlayed in different colors (first frame = red, 30^th^ frame = green). Filaments that had moved, detached or landed during this time span appear either red or green, whereas immobile filaments appear yellow. Those filaments were counted manually and the percentage of total filaments was calculated.

To compare the gliding velocity in the presence or absence of caldendrin, we used the “manual tracking” plug-in of FIJI/ImageJ (*Cordelières F (2005) Manual Tracking, a plug-in for ImageJ software. Institut Curie, Orsay, France*) to track individual actin filaments by one of their ends. Frame-to-frame velocity was calculated from these tracks.

### Statistical Analysis

Statistical analysis was performed in Prism 6.05 (*GraphPad*; all other tests). Detailed specifications about the type of test, significance levels, n numbers, and biological replicates are provided in the figure legends. Experimental repeats and n numbers per experiment were chosen according to experience with effect size. Data are represented as mean ± SEM throughout the manuscript. The data were tested for normality using the D’Agostino-Pearson test (Prism) and accordingly subjected to parametric (t-test, ANOVA) or non-parametric tests (Mann-Whitney test, Kruskal-Wallis test) for significance. The analysis of the caldendrin knock out mice data was done blindly.

## Data Availability Section

Protein interaction AP-MS data are provided in Table AV1.

## Acknowledgments

The authors would like to thank W. Wagner (*GFP-myo Va fl*), T.G. Oertner (*ER-tDimer2*), and L. Kapitein and C. Hoogenraad (*KIF17_1-547_-GFP-PEX26, PEX-mRFP-FKBP, MyoVb_1-1090_-GFP-FRB*) for sharing plasmids. jGCaMP7s was a gift from D. Kim (*Addgene #104463*). We further thank S. Hochmuth for preparing the mouse neuronal cultures, J. Bär and M. Andres-Alonso for preparing the rat neuronal cultures, R. Raman for providing the purified caldendrin Ca^2+^ binding mutant protein, the UKE Microscopy Imaging Facility (UMIF) and the Leibniz-Institut for Neurobiology Magdeburg (LIN) for access and use of their spinning disc and confocal microscopes. This work was supported by the Deutsche Forschungsgemeinschaft (DFG Emmy Noether Programme MI1923/1-2, FOR2419 TP2, SFB877 B12 and Excellence Strategy – EXC-2049–390688087) and Hertie Network of Excellence in Clinical Neuroscience and Excellence Strategy Program.

## Author Contributions

AK and NH cloned expression constructs and performed co-immunoprecipitations, AK performed biochemical and cell biological experiments, DHWD and JAAD performed mass spectrometric analysis, JG performed mass spectrometric, ER tracking and MSD analysis, AK, JG and MM analyzed the results and wrote the manuscript, all authors read the paper, MM acquired the funding and coordinated the project.

## Tables and their legends

**Table EV1. Mass spectrometry of caldendrin pull-down**

The full result of the mass spectrometric analysis of caldendrin pull-down from brain lysate, sorted by gene names. Processing of this data is described in the methods section. Note the predominantly high detection intensities for peptides only in the cald-GFP Ca^2+^ condition, indicating a general lack of binding in its folded state.

## Expanded View Figure legends

**EV Figure 1: Mass-spectrometric analysis of brain-specific caldendrin interactors hints at myosin family as a new class of Ca^2+^-dependent binding partners.**

**A** Coomassie blue stained SDS-gel of purified bio-GFP-caldendrin or bio-GFP (control) and interacting proteins pulled down from mouse brain extract in the presence (Ca^2+^) or absence (EGTA) of calcium. Arrows indicate the bands representing bio-GFP and bio-GFP-caldendrin.

**B** Proteins identified by mass spectrometry in the bio-GFP-cald pulldown from brain lysate displayed as ratio of Ca^2+^/EGTA conditions and corrected for possible non-specific binding to GFP. Myosins are indicated in red. Proteins indicated with * contain an IQ motif, according to their UniProt entry. Also see EV Table 1.

**C** Putative IQ-motif sequences of proteins in **B** were extracted (UniProt), aligned in MATLAB and subsequently visualized in Weblogo (https://weblogo.berkeley.edu/logo.cgi).

**EV Figure 2: Purification of in vitro assay components and F-actin pelleting assay**

**A** Coomassie blue stained SDS-PAGE showing purified full-length myoVa (arrow) from HEK293 cells using a StrepTag-GFP-tag.

**B** Western blot analysis of purified StrepTag-GFP-myoVa. After elution from StrepTactin beads using biotin, the remaining protein was eluted by boiling in SDS-loading buffer. Biotin-induced elution was incomplete, however the remaining, non-elutable fraction of myoVa seems to be not associated with CaM. Endogenous CaM is present in the eluted fraction independent of the availability of calcium during the binding step.

**C** Coomassie blue stained SDS-PAGE showing untagged CaM purified from *E. coli* using hydrophobic interaction chromatography.

**D** Coomassie blue stained SDS-PAGE showing untagged caldendrin (cald) purified from *E. coli* using a “self-cleaving” intein tag.

**EV Figure 3: Verification of the quadruple transfection efficiency in COS7 cells**

**A** COS7 cells were transfected with the calcium indicator GCaMP7s to assess the endogenous calcium signaling activity of the cell line. Time-lapse imaging shows spontaneous calcium spikes over a time course of 5 minutes. Also see Movie EV3.1.

**B** Representative microscopy images of COS7 cells co-transfected with inducible dimerization and peroxisomal labeling constructs (MyoV-GFP-FRB, KIF17-GFP-PEX, PEX-RFP-FKBP) and additionally expressing either untagged caldendrin or control empty vector. Cells were fixed and stained using an anti-caldendrin antibody to show co-expression rate. Scale bar = 100 μm.

**C** Co-immunoprecipitation of caldendrin-tagRFP (cald-tagRFP) with the myoVb construct used in Fig. 3C (MyoVb_1-1090_-GFP-FRB). The western blot shows Ca^2+^-dependent binding of cald-tagRFP to the myoVb construct.

**D** Western blot analysis of F-actin sedimentation assay (triplicate). Full-length GFP-tagged myoVa was co-expressed with cald-tagRFP (or tagRFP only) in HEK293 cells. The cells were lysed in F-actin stabilization buffer, the lysate was supplemented with Ca^2+^ and F-actin was pelleted via ultracentrifugation. The presence of caldendrin had no effect on the binding of myoVa to F-actin.

**EV Figure 5: SER spine entry dynamics upon overexpression of caldendrin**

**A** Visualization of ER presence over time in individual dendritic protrusion of control (GFP) or caldendrin over-expression (cald-GFP) neurons as shown in **Fig 5A**. Red = no ER present, white = ER present.

**B** Quantification of **A**, also see **Fig 5A, B**. No difference in ER entry dynamics in these protrusions can be seen.

**C** Quantification of spine and filopodia density upon overexpression of cald-GFP. Corresponds to **Fig 5C**.

